# Open-source virtual reality platform for head-fixed adult zebrafish

**DOI:** 10.64898/2026.10.19.760147

**Authors:** Alexandre Javier, Sriram Narayanan, Lukas Anneser, Jaap van Krugten, Rainer W. Friedrich

## Abstract

Virtual realities (VRs) are powerful tools to study how brains interact with the world through behavior. We present a new platform for closed-loop high-resolution virtual-reality experiments in head-fixed adult zebrafish. The VR is defined using open-source software, projected onto a semi-hexagonal tank by three projectors, and controlled by software allowing for modular trial-and-block experimental designs. Real time VR updates are performed by an optimized algorithm based on infrared tail videos. Two-photon imaging during behavior is achieved by custom gating of VR projectors and coordinated with stimulus triggers by a microcontroller. Using this system we found that fish respond to visual landmarks in virtual corridors with distinct changes in locomotion parameters and body posture. Adding landmarks to a familiar corridor elicited localized behavioral responses that decayed with experience. Restricting the viewing distance by simulating “murky water” further confirmed that behavioral responses were driven by the visibility of landmarks independent of their position. Our platform provides a user-friendly, versatile VR system for the joint analysis of neuronal activity and behavior in adult zebrafish.

## 1 Introduction

Virtual realities (VRs) are powerful tools to study how brains control behavior because they enable systematic manipulations of the world that are unrealistic in physical environments. Moreover, VRs facilitate activity measurements during behavior and parameterized analyses of neuronal and behavioral responses to sensory input (Heisenberg and Wolf, 1979; Fry et al., 2008; Ahrens et al., 2012; Jouary et al., 2016; Radvansky and Dombeck, 2018). In larval zebrafish, VRs can be combined with brain-wide imaging of neuronal activity and have been used to address a broad range of questions. For example, VR-based approaches provided insights into the mechanisms underlying visuo-motor reflexes (Orger et al., 2008; Ahrens et al., 2013), prey capture (Bianco et al., 2011), the representation of space (Yang et al., 2024), and the integration of information over time (Yang et al., 2022). However, social behavior and various forms of learning emerge only at later developmental stages, indicating that the cognitive behavioral repertoire of larval zebrafish is limited. Adult zebrafish, in contrast, show relational and configural learning, reversal learning, and delayed matching-to-sample performance, which are only rudimentary in larvae (Gerlai, 2017; Bloch et al., 2019). Moreover, adult zebrafish show social learning, observational learning and the ability to learn abstract rules (Lindeyer and Reader, 2010; Oliveira, 2013; Chou et al., 2016), indicating that cognitive abilities are well-developed (Friedrich et al., 2010; Oliveira, 2013). This notion is further supported by the ability of other teleosts to learn abstract structures and rules (Rodríguez et al., 2021; Schluessel et al., 2022). While optical accessibility of the brain in adult zebrafish is limited compared to larvae, neuronal activity in the dorsal telencephalon can still be measured by multiphoton calcium imaging through the intact skull. Recently, activity related to prediction, navigation and learning has been revealed in the telencephalon of adult zebrafish by imaging neuronal activity in a VR under head fixation (Huang et al., 2020; Torigoe et al., 2021; Islam et al., 2024; Palacios-Flores et al., 2026). Nonetheless, VR approaches in adult zebrafish have not yet been widely adopted, possibly because experimental procedures remain complex. We therefore developed a technical platform and streamlined procedures to facilitate VR experiments.

The success of VR experiments depends on multiple factors including the following. (1) Many VR experiments include head fixation to enable optical activity measurements with stationary microscopes. While larval and young juvenile zebrafish can be held in place by embedding the head in agarose, older zebrafish require more stable fixation, which is achieved by attachments glued to the skull (Huang et al., 2020; Torigoe et al., 2021; Bandonil et al., 2023; Islam et al., 2024). One important requirement is therefore a stable head fixation method that does not obscure optical access to the brain area of interest. (2) Closed-loop VR updates under head fixation are typically performed by predicting swimming trajectories based on tail videos in real time. We observed that prolonged naturalistic bout-and-glide swimming of adult zebrafish required a closed-loop configuration with efficient VR updates (Huang et al., 2020), indicating that the behavioral state is sensitive to the accuracy of trajectory predictions. The algorithm for trajectory predictions and the implementation of closed-loop updates are therefore critical components of a VR system. (3) Many experiments include flexible variations of a VR, such as the addition or removal of visual objects on a trial-by-trial basis or in response to behavioral actions. VR systems should therefore offer options for interactive manipulations. (4) As technical complexity can be a limiting factor the implementation of VR systems should be streamlined as much as possible. We specifically considered these factors in the design of our VR platform. Moreover, we included an optional perfusion system to maintain constant water flow over the fish and to apply odor stimuli.

In VRs for zebrafish, tail movements are usually analyzed to update the VR by predicting linear and angular acceleration or displacement. In principle, however, the underlying video data contain additional information about body posture. We therefore explored whether a more fine-grained analysis of video data can provide additional information about behavioral states and behavioral engagement. In rodents, activity of hippocampal place cells critically depends on active exploration (O’Keefe and Dostrovsky, 1971; Harvey et al., 2009) and spatial coding in a VR deteriorated substantially when mice disengaged from a spatial task (Kentros et al., 2004; Pettit et al., 2022; Ortiz et al., 2025). We therefore specifically analyzed swimming behavior that may be indicative of behavioral engagement with the VR.

We present a platform for VR experiments in adult zebrafish that includes procedures for stable head fixation, a high-resolution panoramic projection system, a two-photon microscope for minimally invasive imaging of telencephalic activity, a system for water perfusion and odor stimulation, technical solutions to integrate all system components, and software to design flexible and modular experimental schedules. The system was used to analyze the behavior of adult zebrafish traversing virtual corridors with naturalistic textures and distinct visual objects (landmarks). To examine responses to landmarks independent of position we simulated “murky water” by gradually decreasing visibility as a function of distance relative to the fish. While fish typically showed naturalistic swimming patterns in the VR throughout an experiment, behavioral engagement, as assessed based on activity measures, was found to be variable, possibly reflecting variations in brain states and motivation. We further observed position-independent behavioral responses to novel landmarks. Consistent with previous observations, neurons in telencephalic area cDc showed activity at specific locations in the VR. Our VR platform is therefore a versatile tool to study neuronal activity, visually guided behavior, and navigation in adult zebrafish.

## 2 Results

### 2.1 Head fixation and system overview

Our VR platform is based on the system described by Huang et al. (Huang et al., 2020). While this system functions reliably, as assessed by the maintenance of naturalistic motor behavior over extended periods of time, its implementation is demanding. A main goal was therefore to simplify experimental procedures and system components to facilitate VR applications in practice.

Head fixation was originally performed by gluing two minature stainless steel pins to specific bones on both sides of the skull (Huang et al., 2020). Subsequently, stainless steel pins were replaced by headplates, which facilitated and accelerated surgical procedures (Bandonil et al., 2023) and provides options to position additional devices relative to the fish (Torigoe et al., 2021; Islam et al., 2024). We made minor modifications to the headplate design of Bandonil et al. (Bandonil et al., 2023), which consists of a central bar and two C-hooks that are mounted by screws to standard stainless steel posts. To minimize clipping of the effective numerical aperture (NA) of an objective over the telencephalon we modified the shape of the headplate to position the head of the fish further away from the C-hooks and posts (Fig. 1a-c). The headplate was glued to the cranium posterior to the telencephalon using dental cement, which afforded high stability with minimal obstruction of the telencephalon.

**Figure 1.**
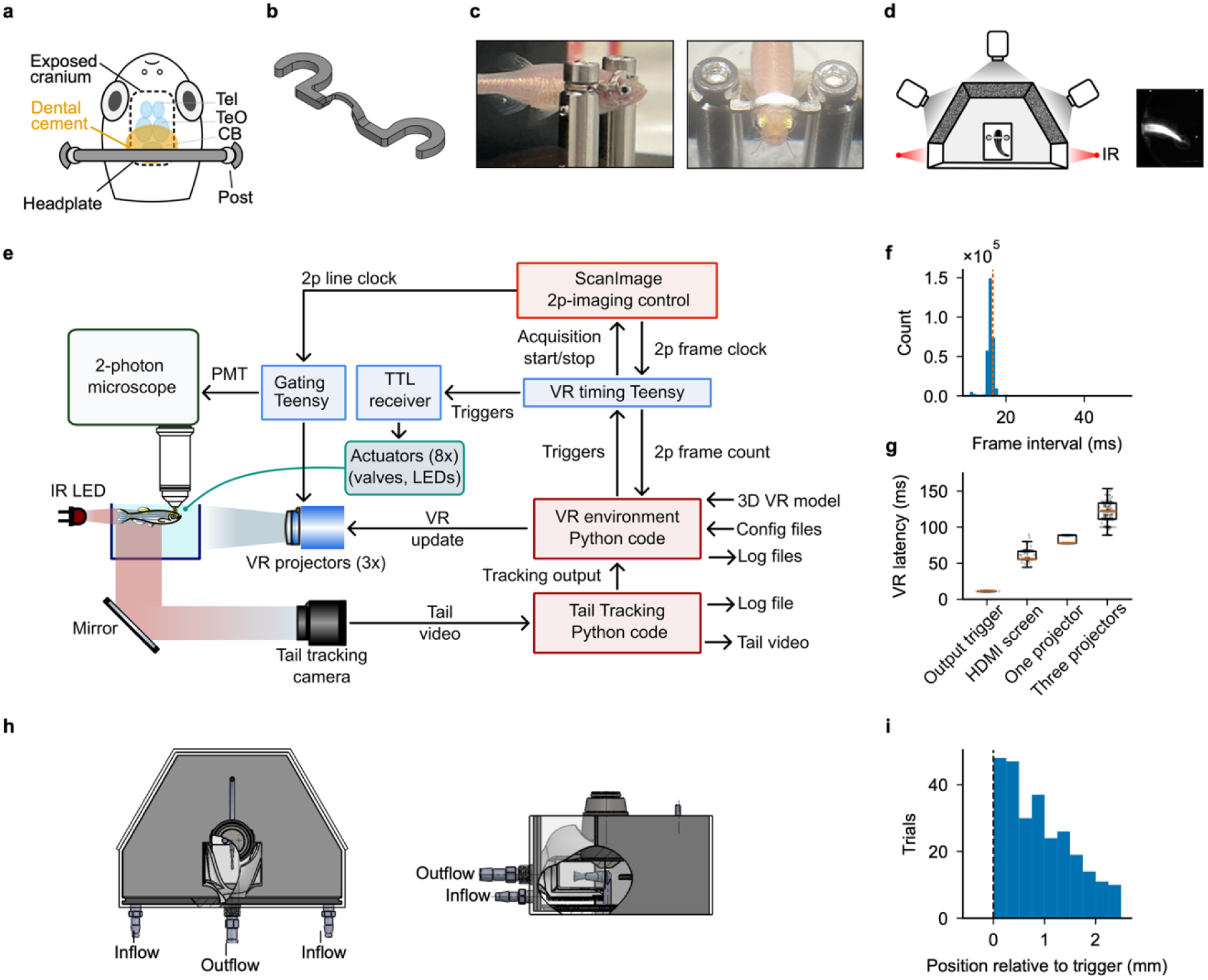
Virtual reality for head-fixed adult zebrafish. **a**, Schematic: head-fixation, dorsal view. The cranium is exposed over the telencephalon (Tel), optic tectum (TeO) and cerebellum (CB); the headplate is bonded to the skull with dental cement caudal to the window and fixed to the posts by C-hooks. **b**, Headplate. **c**, Fish with headplate secured to the posts. **d**, Top view of the tank with infrared (IR) illumination. The VR is projected onto the semi-hexagonal walls of the tank by three projectors, covering 180°. Right: image of the tail on the IR camera from below under dark-field illumination. **e**, Hardware/software architecture of the closed loop VR. The projection is updated by tracking tail motion in IR video frames in real time. A timing microcontroller (Teensy) synchronizes stimulus triggers, actuators and the two-photon acquisition. **f**, Distribution of VR frame intervals (n = 3.10 × 10^5^ intervals, 0.8 ms bins). The mode is 16.0 ms and 92.5% of intervals fall between 14.8 and 17.2 ms. Dashed line marks the 16.67 ms period of the 60 Hz configuration. **g**, VR update latency measured directly at the level of the output trigger (n = 448) and as changes in the display on an HDMI screen (n = 211) or projections by one projector (n = 296) or three projectors (n = 259). Medians 11.11, 55.64, 77.78 and 122.22 ms; means ± s.d. 11.11 ± 0.00, 60.35 ± 6.56, 82.62 ± 5.51 and 120.33 ± 12.91 ms. Boxes show median and interquartile range; whiskers show 1.5 × interquartile range; points show individual measurements. **h**, Fish tank with perfusion and odor delivery system, top and side views. **i**, Spatial precision of behavior-dependent trigger. Histogram shows the distribution of the position of fish in a corridor (1.6 m length) relative to the defined trigger position (n = 276 trials, 3 sessions, 2 fish).

Mounting posts were fixed to a baseplate fitting into a matching hole in the bottom of the holding tank. The fish head was placed approximately in the center of the holding tank, which consists of three walls arranged in a semi-hexagonal pattern and a short extension in the caudal direction (Fig. 1d) (Huang et al., 2020). The VR was projected onto a diffusive film lining the outside of the three walls by three projectors. The virtual world was created in Blender and rendered in Panda3D as a single 5,760 × 1,080 pixel window split into three display regions, one per projector. Regions were drawn by three virtual cameras, each covering a 60° sector. The visual scene therefore extended for ∼180° around the fish (azimuth) with a height (elevation) of ∼34° (Fig. 1d). Regions behind the fish were invisible. Video images of the tail (Fig. 1d, right) were acquired from below through the transparent floor under dark-field infrared illumination that is invisible to the fish.

VR updates require the conversion of each tail video frame into a forward velocity and a heading estimate for transmission to the VR engine. Because previous procedures were complex and inflexible we implemented this task *de novo* (Fig. 1e). The VR engine, written in Python on Panda3D, received information asynchronously via the user datagram protocol (UDP) to update the scene and to drive the three projectors. A microcontroller (Teensy 4.0) running custom firmware was used to generate and receive digital triggers required for synchronizing external processes such as two-photon acquisition and stimulus delivery with the VR experiment. The microcontroller issues start and end triggers for two-photon image acquisition, reads the two-photon frame clock and acquisition state back from the microscope hardware, and carries eight stimulation trigger outputs plus one analogue intensity line for actuators such as optional olfactory stimulation valves and LEDs. Trigger pulses requested to be executed on a particular VR frame are cached and executed in the subsequent frame in case of synchronization errors, which may arise due to the asynchronous nature of the USB interface. Each process writes its own time-stamped log file and the tail video is stored alongside the tracking log. Any session can therefore be re-analysed from the raw frames. The modular nature of the tail-tracking workflow, the VR rendering loop and the microcontroller interface, together with the asynchronous data transfer between these processes, greatly facilitates changes in system components such as projector configurations and camera hardware.

Performance data were read directly from the log files and showed that the VR frame interval was narrowly distributed. In a typical set-up running at 60 Hz, the modal interval was 16.0 ms and 92.5% of 3.10 × 10^5^ intervals fell between 14.8 and 17.2 ms (Fig. 1f). We further measured the lags between an update command, the corresponding output trigger signal (∼11 ms), and its detectability as changes in the display on an HDMI screen (∼56 ms) or projections using one or three projectors (∼77 ms and ∼122 ms, respectively; Fig. 1g). Changes in the visual output were measured directly using a photodiode. These results are consistent with VR update lags in other VR systems (Stowers et al., 2017) and confirm that the effective lag is dominated by the graphics components.

In previous VR systems, fish were held in static water during the experiment. To add an option for water perfusion and odor stimulation we added two inflow ports on the rear side of the tank near the lateral walls and an outflow port attached to a 3D-printed funnel extending around the trunk of the fish (Fig. 1h). The lateral walls of the funnel were made of transparent acrylic to allow for illumination and observation of the tail. This arrangement resulted in water flow extending from the rear towards the front of the holding tank, along the fish in rostro-caudal direction, and into the funnel. In addition, a narrow odor delivery tube held by a micromanipulator outside the tank could be placed in front of the fish. Odor stimuli therefore targeted the nose and subsequently entered the exhaust. Dye tracing confirmed that pulses of liquid applied through the odor delivery tube were collected effectively by the funnel without dissipating into the tank.

Olfactory or other external stimuli should usually be triggered when the fish reaches a defined location in the VR. The temporal precision of stimulation depends on multiple factors including the timing of closed-loop and trigger signals and the resolution of VR updates. To measure this precision we defined a trigger at a specific location along a 1.6 m long linear virtual corridor and measured the actual position of a fish at the time of the trigger signal. On average, the fish was located 0.88 ± 0.75 mm ahead of the specified location at the time of trigger (276 trials from 2 fish; Fig. 1i). Hence, the spatial inaccuracy of the trigger signal was minimal, probably reflecting the delay in the VR update and confirming high spatio-temporal resolution of the VR.

### 2.2 Conversion of tail kinematics into virtual locomotion for closed-loop VR updates

A key component of the VR system is the procedure that translates tail movement into the locomotor signal driving VR updates. This procedure runs as a standalone application that acquires frames from the camera, tracks the tail in real time, computes velocity and heading values, displays them to the operator, writes them to a log together with the raw video, and sends them to the VR engine (Fig. 2a). Acquisition and tracking parameters are read from, and written back to, a setup file to store the exact parameters for each session. The procedure described by Huang et al. was designed for one specific camera model that is no longer available (Huang et al., 2020). We therefore updated the software to support two commonly used camera families.

**Figure 2.**
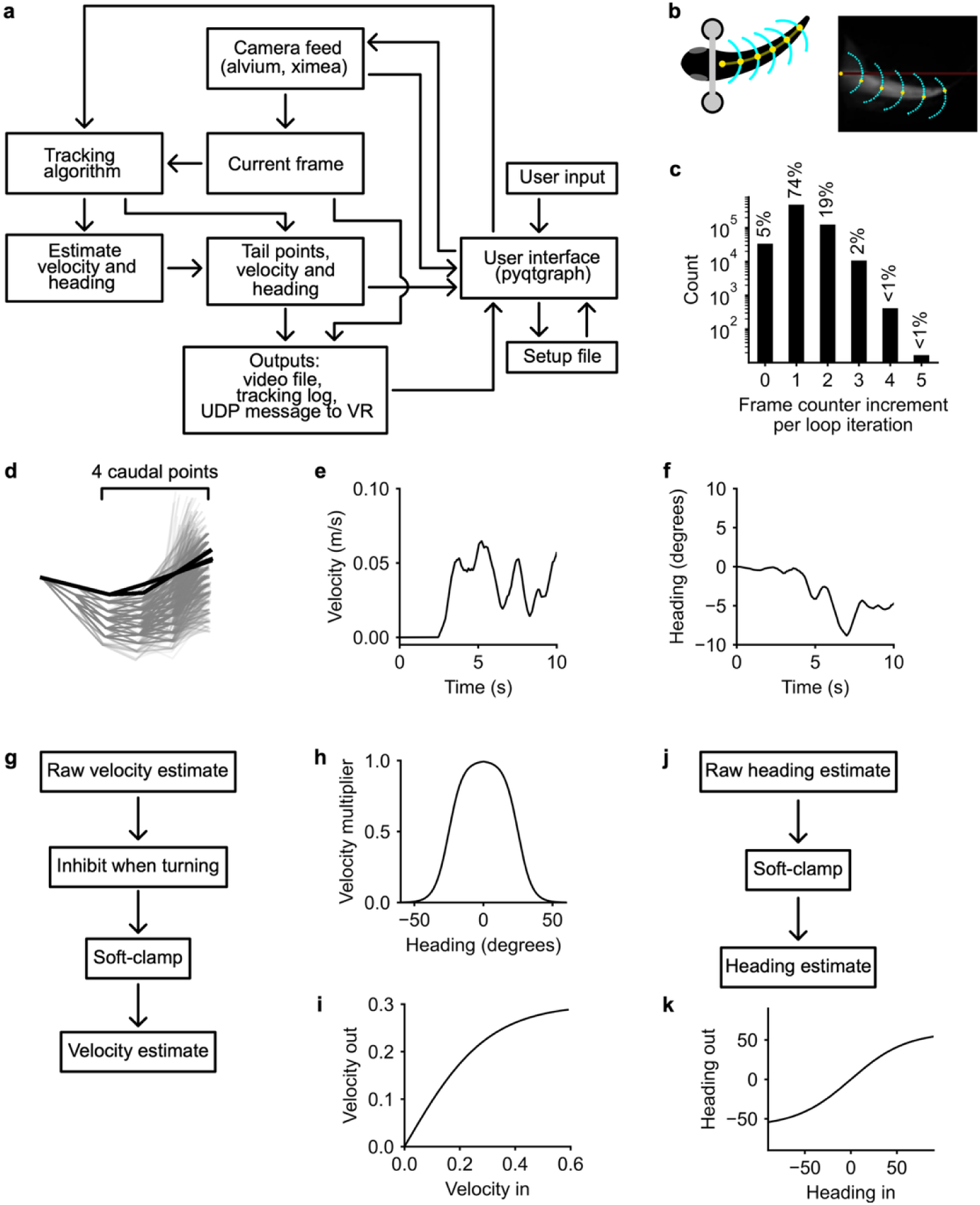
Tail tracking and velocity/heading estimators. **a**, Architecture of tail tracking software. **b**, Schematic: fixed start point at the base of the tail and the chain of tracked points. Right, camera frame with the tracked points (yellow) and the search arcs (cyan). Red line: reference axis. **c**, Distribution of the camera frame-counter increment between successive iterations of the tracking loop (a direct measure of how the tracker keeps up with the camera). An increment of 1 means that every camera frame was tracked; 0 means that the loop ran twice on the same frame; >1 means that frames were skipped. **d**, Tail shapes from one swim event. **e, f,** Example of forward velocity (e) and heading (f) returned by the tracker over 10 s. **g**, Processing chain for forward velocity. Over the 0.4 s estimator window (60 frames at 150 Hz), the offset-corrected cumulative tail angle is split into its positive and negative parts. The raw estimate is twice the smaller of the two summed magnitudes, scaled by the forward gain, so that only alternating tail movement produces forward motion. The estimate is then attenuated as a function of heading and soft-clamped. **h**, Turn-inhibition weight as a function of heading. With the parameters used in this study (logistic midpoint 15, steepness 20) the weight was nearly constant, varying between 0.59 (straight tail) and 0.52 (largest heading observed). **i**, Velocity soft-clamp, y = v_max_ tanh(x/v_max_). In the sessions reported here v_max_ = 0.30 m s_−1_. **j**, Processing chain for heading. **k**, Heading soft-clamp. With the parameter used in this study the clamp was nearly linear over the entire operating range.

We redesigned procedures for VR updates to facilitate their implentation and to further improve the quality of locomotor predicitions from tail movements. The tail tracking algorithm is iterative and needs no model of the fish (Fig. 2b). From a fixed start point at the base of the tail, the tracker moves along the tail in steps of 30 pixels. At each step it takes the brightest pixel on an arc of ±1.2 rad (0.12 rad steps), centered on the direction of the preceding segment. Six such steps give seven tail points per frame. The angles between consecutive points are the raw signal. The sum of the last four segment angles is defined as the cumulative tail angle, which enters the estimators. A running median of the cumulative tail angle over the preceding 15 s is subtracted as an adaptive offset to ensure that a slow drift in resting posture is not interpreted as a sustained turn.

To determine whether the tail tracking procedure keeps up with the camera we measured the increment of the camera’s hardware frame counter between successive iterations of the tracking loop (Fig. 2c). An increment of 1 means every frame was tracked, 0 means that the loop ran twice on the same frame, and 2 or more means that frames were skipped. In the example recording used, the tracker kept up on 74% of iterations and skipped a frame on 19%. Because the velocity and heading estimators sum the cumulative tail angle over a 0.4 s window (60 frames at 150 Hz), an occasional skipped frame removes one sample in sixty and has a negligible effect on the locomotor signal. However, it is important that skipped frames are logged to ensure temporal precision in the analysis, as performed by our system.

During movement of the fish, the tracked tail shapes fanned out around the resting posture, with the four caudal points moving the most (Fig. 2d). The two outputs of the tracking algorithms are forward velocity and heading direction. As shown in the example in Fig. 2e,f, forward velocity typically rose from zero at the onset of a swim bout and decayed between bouts, while heading fluctuated around zero, sometimes showing a sustained lateral bias.

Forward velocity is computed so that only alternating bilateral tail movement drives the fish forward (Fig. 2g). Over the 0.4 s time window of the estimator, the offset-corrected cumulative tail angle is split into its positive and negative parts. The raw velocity is then calculated as twice the smaller of the two summed magnitudes, scaled by a forward gain. A one-sided bend, however large, therefore produces no forward motion, which removes the most common artifacts. In the latest version of the algorithm, forward drive is weighted down by turn amplitude using a logistic function. The magnitude of the ongoing turn signal is mapped onto a predetermined weight function and the resultant weight is multiplied with the raw forward drive estimate (Fig. 2h). The resultant velocity estimate is soft-clamped as v_max_ tanh(v/v_max_) with v_max_ = 0.30 m s^−1^ (Fig. 2i).

The heading command follows a parallel chain (Fig. 2j). The offset-corrected cumulative tail angle is summed over the same 0.4 s window, scaled by a heading gain, soft-clamped in the same tanh form (Fig. 2k), low-pass filtered with a 400 ms time constant and zeroed inside a small dead-zone. In the sessions reported here the heading clamp parameter was 15 rad (859 degrees). Note that in the linear corridors used in this study, the heading signal had no influence on VR updates but it was nonetheless used as a behavioral read-out of lateralized tail activity. The heading gain was identical in all 25 “Landmark addition” and “Landmark deletion” sessions (see below), allowing for quantitative comparisons of cumulative tail angles between animals.

### 2.3 Trial editor and modular design of experiments

An open-source software environment was established for the modular design of experimental schedules (Fig. 3; see Methods for software availability). Each trial is defined by a families of parameters: trial structure and locomotion (start and end coordinates, distance threshold, maximum duration, inter-trial interval, gains, fade), stimulation channels (up to eight, with per-channel coordinates, radius, crossing direction, onset and duration), optic-flow perturbations, landmarks (number, positions, orientation, scale and texture) and collisions (Fig. 3). These parameters are set through a user interface, the Trial Editor. Trials are assembled into blocks with a repeat count and optional randomization (Fig. 3, 6), and blocks are then chained into an experiment for execution by the VR engine. The full per-trial configuration is written into the session metadata.

**Figure 3.**
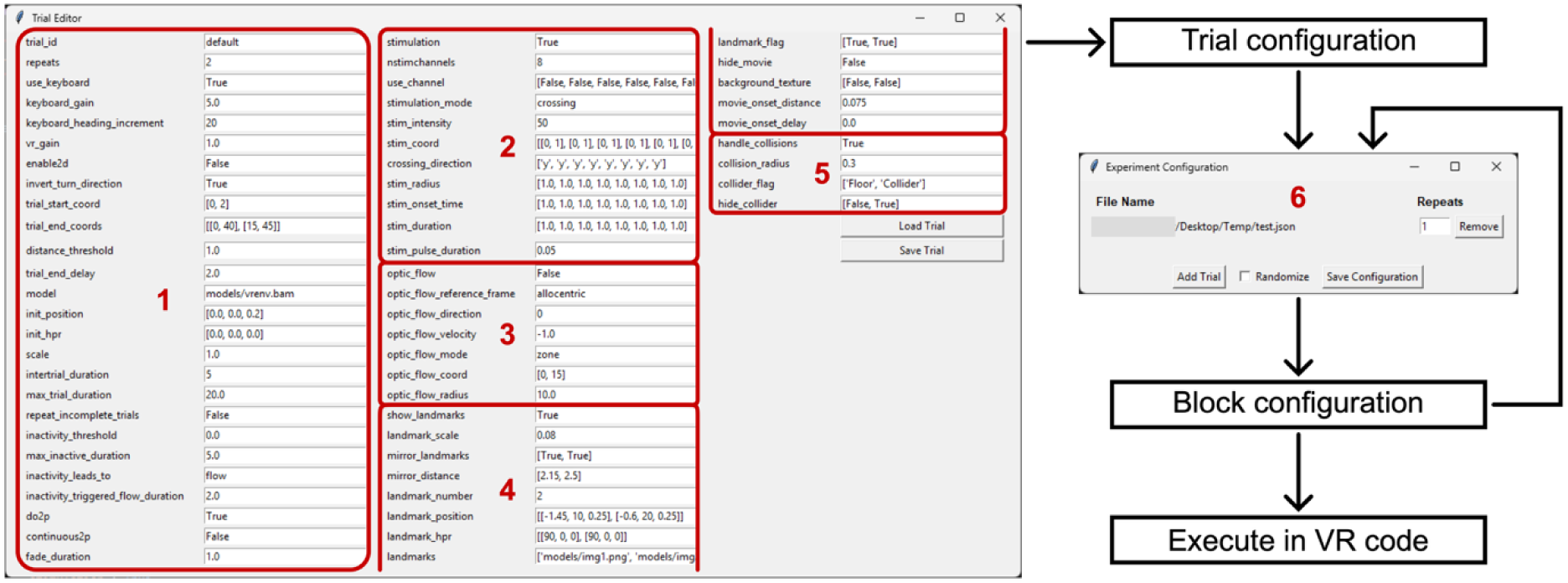
User interface for experimental design. Left: Trial Editor main window. Parameters of one trial are displayed in groups: (1) trial structure and locomotion, (2) stimulation channels, (3) optic-flow perturbations, (4) landmarks and (5) collisions. Saved trials are assembled into blocks with a repeat count and optional randomisation in the experiment configuration window (6). Right: chain from trial configuration to block configuration to execution in the VR engine. The full per-trial configuration is written into each session’s metadata.

### 2.4 Analysis of swimming patterns

We examined the behavior of adult zebrafish in a 1.6 m long virtual corridor with naturalistic textures and optional landmarks (n = 34 fish). When the fish reached the end of the corridor, lights were briefly dimmed and the fish was teleported back to the start (Fig. 4a). One trial corresponded to one corridor traversal and one session corresponded to a continuous recording of a single fish. A session therefore comprised multiple trails, potentially using different VRs and parameters, that may be organized into blocks. Data from each session is logged as one set of files. Sessions lasted from ∼40 min to >3 h and some fish were recorded in more than one session. In most experiments, corridors contained a single visual landmark that was present or absent in different trials. The landmark was a stone arch with an aperture substantially larger than the fish. As exploratory observations in freely swimming fish indicated that adult zebrafish avoid narrow apertures, we surmised that an arch of moderate size may have subtle effects on behavior without disrupting continuous swimming.

**Figure 4.**
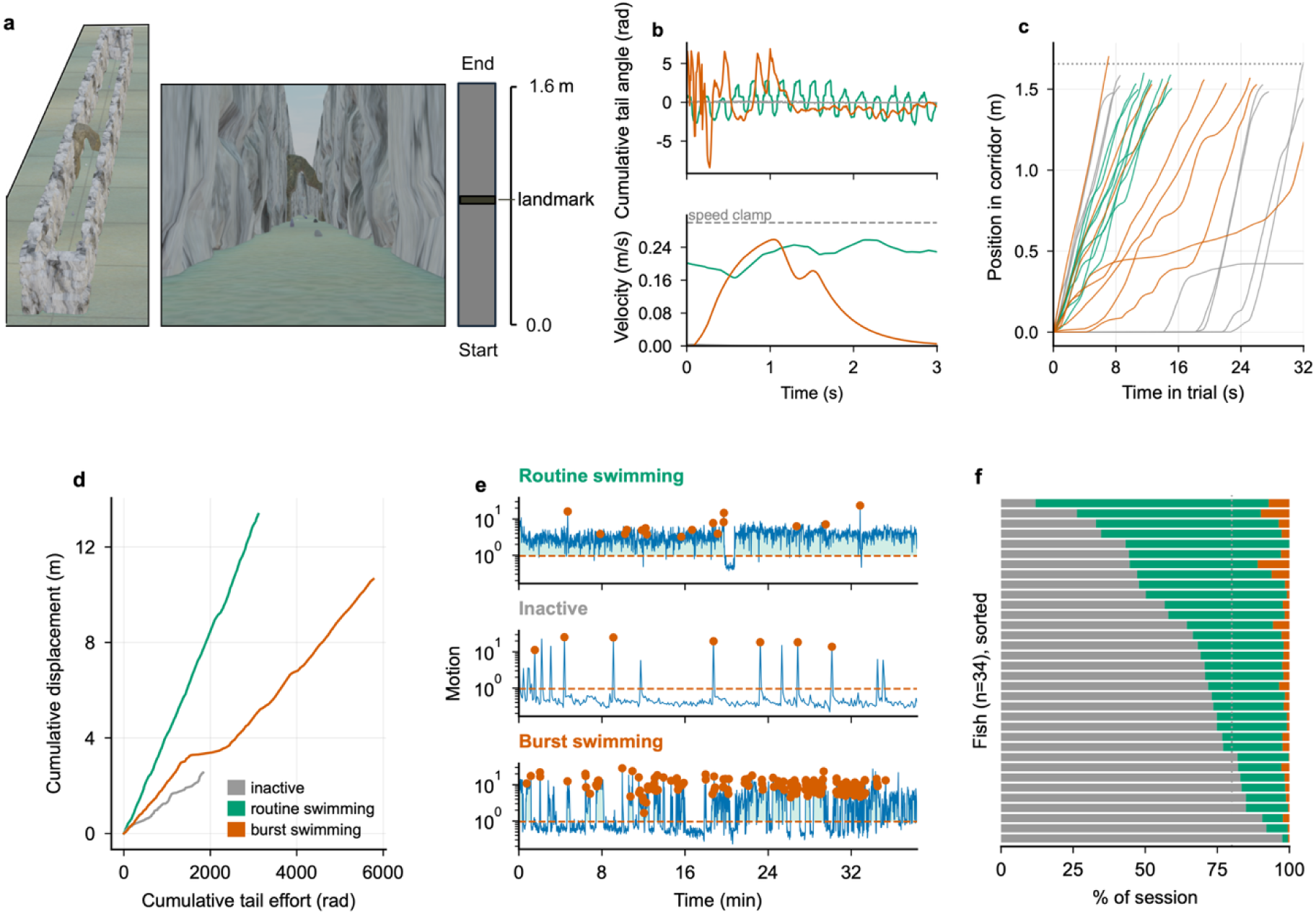
Classification of behavioral state. **a**, Virtual corridor. Left: top & front view into the corridor at the start. Right: top view and schematic layout. **b**, Cumulative tail angle (top; mean-centered) and swimming velocity (bottom; dashed line marks speed clamp at 0.30 m/s) of three fish (different colors) during 3 s time windows. Fish were chosen to represent different behavioural classes (routine swimming, burst swimming, inactive; see color code in d). **c**, Position along the corridor as a function of time within a trial. Dotted line marks the end of the corridor (1.6 m). Each trace corresponds to one trial. **d**, Cumulative displacement as a function of cumulative tail effort for the same three fish integrated over one session. **e,** Motion of the same three fish as a function of time (first 38 min of a session). Green shading, epochs classified as active; dashed line, activity threshold; orange points, windows classified as burst swimming. **f,** Classification of behavioral states across all fish (n = 34), sorted by engagement. Each horizontal bar shows the fraction of time spent by one fish spent in each of the three behavioral states.

Fish showed different swimming patterns that were assigned to one of three behavioral states (Fig. 4b-d): (1) Routing swimming. This pattern was characterized by extended periods of continuous low-amplitude swimming, typically showing a bout-and-glide pattern. (2) Burst swimming. This pattern consisted of intermittent high-intensity tail motion, including c-bends or struggles, separated by periods of no swimming. (3) Inactivity. In this state, fish were stationary most of the time and only occasionally showed brief epochs of locomotion.

To assess swimming behavior more systematically we quantified two parameters: (1) a motion index based on frame-by-frame intensity changes of bright pixels, and (2) a tail bend index based on an aspect ratio estimate of the tail shape (Methods). In each video frame, fish were defined as inactive when the activity index was below a pre-defined threshold. Otherwise, the fish was defined as active if the tail bend index was below a threshold or as burst-swimming otherwise (Methods). The fraction of time spent in in the active or inactive state varied widely between individuals while burst-swimming was rare in all fish (Fig. 4f). Within each session, a fish was classified as inactive if it spent <20% of the time in the active state (n = 9 fish) and as burst-swimming if it spent >5% of the time in the burst-swimming state (n = 5). Otherwise, fish were classified as swimming routinely (n = 20).

Routinely swimming fish traversed the corridor most effciently, frequently reaching velocities near the 0.30 m s−1 speed clamp limit (Fig. 4b). Fish in the burst swimming state moved, on average, more slowly while inactive fish were the slowest. To further examine swimming efficiency we estimated “tail effort” by summing the local bend angles along the tail skeleton within each frame. The results indicate that routinely swimming fish generated displacement with the least effort, followed by burst-swimming and inactive fish (Fig. 4d). Burst swimming was less efficient than routine swimming because the tail bends were often unilateral, resulting in little or no forward displacement.

### 2.5 Behavioral responses to visual landmarks

We next examined whether swimming behaviour was influenced by the visual landmark (arch) using two experimental configurations: (1) “Landmark addition”: The landmark was added after fish were familiarized to the corridor in the absence of the landmark (13 sessions from 10 fish). (2) “Landmark deletion”: The landmark was removed after familiarization in the presence of the landmark (9 sessions from 8 fish). Experiments were structured into blocks of 10 trials (Fig. 5a). Familiarization consisted of 5 or 8 blocks without or with landmark, followed by one block of the opposite type. Thereafter, fish were exposed to pseudo-random sequences of blocks with or without landmark. Visual observations indicated that body posture changed as fish approached the landmark. To quantify changes in behavior we therefore analyzed three readouts of locomotion and body posture: (1) swimming speed, (2) the cumulative tail angle, and (3) “tail complexity”, which was defined as the standard deviation of the local bend angles along the tail skeleton in each frame.

**Figure 5.**
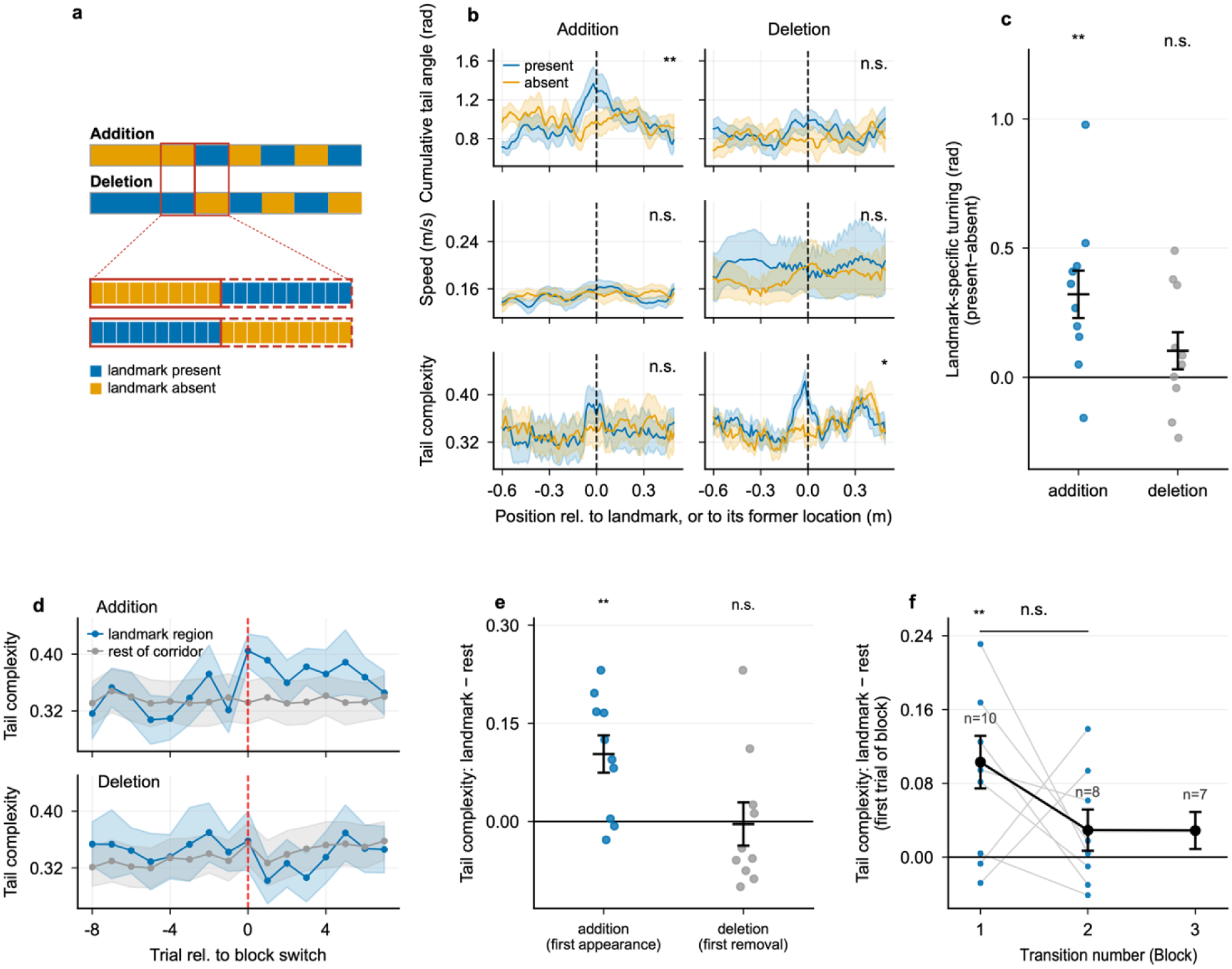
Behavioral responses to a visual landmark (stone arch). **a**, Landmark addition and deletion paradigms. Top: block structure of a session. Sessions always started with at least five blocks (5 × 10 trials) of the the same type (presence or absence of landmark). Thereafter, the block type was switched, and blocks continued to change in a pseudo-random sequence. Bottom: trial structure within blocks. **b**, Cumulative tail angle (top), swimming speed (middle) and tail complexity (bottom) as a function of position relative to the landmark (dashed line), averaged over 10 completed trials (1 block) immediatedly before and after the landmark switch. Left: Landmark addition paradigm (n = 10 fish); right: Landmark deletion (n = 8 fish). **c**, Change in cumulative tail angle near the landmark in the Landmark addition and deletion paradigms. The landmark region was defined as the spatial bin (width, 11.7 mm) containing the landmark ± 2 bins. The remainder of the corridor (total length, 1.6 m) contained all remaining bins. Each dot represents one fish; bars show mean ± s.e.m. (one-sample two-sided t-test; addition, p = 0.009, n = 10; deletion, p = 0.21, n = 8). **d**, Tail complexity in the landmark region and the remainder of the corridor in successive trials around the block switch (dashed red line). **e**, Change in tail complexity within the landmark region (landmark region minus remainder of the corridor) in the first trial after the landmark switch. Each dot represents one fish. The response was significantly different from zero in the addition (**, p = 0.006) but not in the deletion paradigm. **f**, Tail complexity response (landmark region minus rest of the corridor) on the first trial of each block after addition of a landmark (Landmark addition paradigm). Transition numbers refer to the first, second and third block switch. Black symbols, mean ± s.e.m. across fish; n, number of fish. Grey lines join datapoints from 8 fish undergoing at least two transitions (n.s., paired two-sided t-test, p = 0.28). The change in tail complexity was significantly greater than zero in the first (p < 0.01) but not in subsequent transitions.

We first analyzed changes in behavioral parameters around the landmark. In the “Landmark addition” configuration, the cumulative tail angle increased significantly shortly before the fish reached the landmark. No obvious change in cumulative tail angle was observed at the corresponding position in the preceding block when the landmark was absent (Fig. 5b,c). In the “Landmark deletion” configuration, only a small change in cumulative tail angle was observed around the landmark that was not statistically significant (Fig. 5b,c). Note that in this configuration, the fish had traversed the VR in the presence of the landmark for 40 or 70 trials (4 or 7 blocks) before the behavior was measured. Tail complexity also exhibited changes near the landmark that were statistically significant in the “Landmark deletion” configuration (Fig. 5b). Swimming speed, in contrast, remained almost unchanged. Hence, fish showed postural changes near the landmark without an obvious change in locomotion.

Further analyses focused on tail complexity in the region around the landmark on successive trials. In the “Landmark addition” configuration, tail complexity increased abruptly after addition of the landmark in 8/10 fish (Fig. 5d,e). The increase in tail complexity in the first trial after addition of the landmark was most pronounced and statistically significant in the first block switch. Subsequently, the increase in tail complexity upon appearance of the landmark decreased with additional block switches (Fig. 5f). In the “Landmark deletion” configuration, removal of the landmark was associated with a small, non-significant decrease in tail complexity (Fig. 5d,e). Together, these results reveal spatially localized behavioral responses to the landmark that appeared to be modulated by novelty.

To further assess whether the behavioral response to the landmark is modulated by a positional memory we limited the visibility to a maximum of 0.5 m in front of the fish, simulating “murky water”. This was achieved by gradually decreasing contrast starting at 0.3 m and reaching zero contrast at 0.5 m relative to the position of the fish (Fig. 6a). The landmark was placed at one of three positions (0.63, 0.99, and 1.35 m) that were not visible at the start. Landmark positions varied pseudo-randomly between blocks (19 sessions from 12 fish; Fig. 6a).

**Figure 6.**
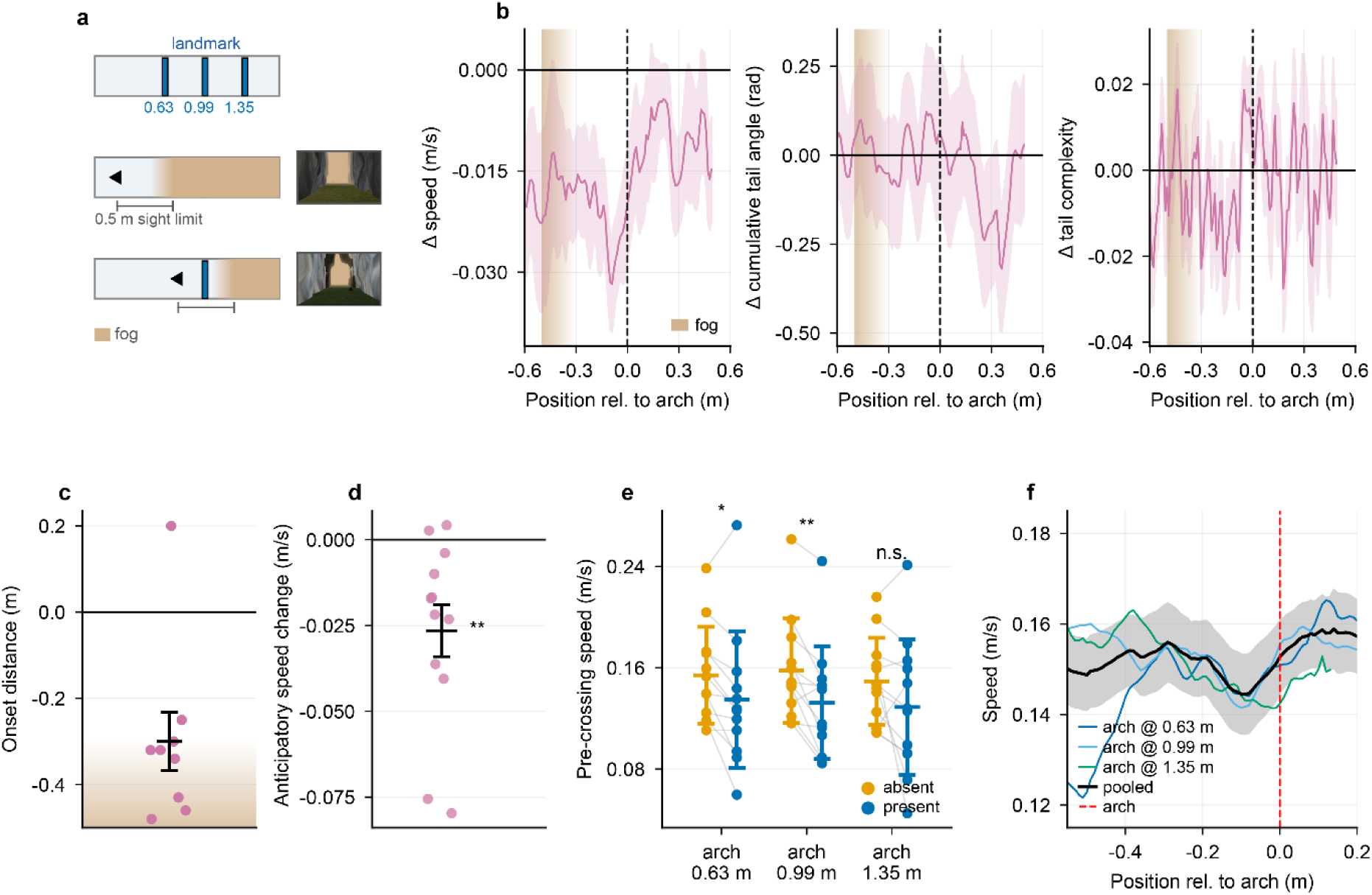
Behavioral responses to landmarks appearing at different positions. **a**, Behavioral paradigm using variable landmark positions in “murky water”. The landmark was located at one of three positions (0.63, 0.99, 1.35 m) along the 1.6 m corridor. Landmark position varied between trials. Visibility decreased with distance from the fish, starting at 0.3 m and reaching zero visibility at 0.5 m (shading). The two corridor views show the same arch at different distances, 0.79 m (invisible) and 0.24 m (fully visible). **b**, Present-minus-absent difference (Δ) in swimming speed, cumulative tail angle and tail complexity as a function of position relative to the landmark (mean ± s.e.m. across fish, n = 12). Shading shows visibility gradient. Between −0.3 and 0 m, speed decreased (−0.022 m s⁻¹, p = 0.01) whereas cumulative tail angle (p = 0.94) and tail complexity (p = 0.51) did not change significantly (one-sample t-tests). **c**, Onset distance of the speed change for each fish (9 of 12 fish with a detectable onset; bar, mean ± s.e.m.). Shading shows visibility gradient. **d**, Pre-landmark speed change in each fish, defined as the mean speed difference between a 1 s time window preceding the crossing of the arch and baseline. Speed was significantly reduced prior to reaching the landmark (mean ± s.e.m; p = 0.007, one-sample two-sided t-test, n = 12 fish). **e,** Mean swimming speed in the 1 s window preceding the crossing at each of the three potential landmark positions in the presence and absence of the landmark (same trials). Lines connect dots representing the same fish; bars show mean ± s.d.. Asterisks depict significance determined by paired two-sided t-tests (0.63 m, p = 0.02; 0.99 m, p = 0.002; 1.35 m, p = 0.11, not significant; n = 12 fish). **f**, Mean swimming speed as a function of position relative to the landmark, pooled separately for each of the three landmark positions. Shading shows s.e.m.

We found that fish reduced their swimming speed when the landmark became visible without obvious changes in cumulative tail angle or tail complexity. Hence, the appearance of the landmark out of “murky water” primarily modulated locomotion rather than body posture. The change in swimming speed became statistically significant (p < 0.05) 0.32 m before the landmark. Consistent with this observation, the first of three consecutive positional bins (10 cm wide) that deviated by >2 standard deviations from the baseline speed was located 0.30 m from the landmark. The magnitude and spatial extent of the speed change, as well as their spatial alignment to the landmark, were similar at all landmark positions (Fig. 6c-e). The appearance of the landmark therefore modulated swimming behavior independent of position, indicating that the behavioral response is driven primarily by visual input. These experiments provide an example of the versatility of the VR platform to implement complex experimental schedules, visual manipulations and behavioral readouts.

### 2.6 Two-photon calcium imaging during virtual navigation

To perform multiphoton calcium imaging during behavior in the VR we gated photomultipliers and projectors using the line clock signal from the resonant scanner. Gating was performed using a dedicated microcontroller (Teensy 4.0) running custom firmware included in the VR codebase (Methods). The photomultiplier tube was gated “on” during line acquisition and “off” during the turnaround phase for 15 µs. The white LEDs of the VR projectors were controlled externally using a custom designed fast voltage-controlled current source (one per projector). The projectors were gated “on” within the 15 µs turnaround window, with a delay of 1 µs after the photomultiplier was gated “off”. The precise duration of the “on” state was determined empirically by reducing the level of bleed-through from residual emission, presumably from the phosphor coating on the projector LED, and was between 4 µs and 11 µs. While the microscope was not scanning the projectors were gated with an 8 kHz pulse train with the same duty cycle as during scans to maintain a constant brightness of the projected scene.

To confirm the functionality of the new VR platform and gating procedures we measured activity in Tg(neurod:GCaMP6f) fish, which express the calcium indicator GCaMP6f throughout the dorsal telencephalon (Rupprecht et al., 2016). Behavioral and multiphoton imaging data were coordinated via the microcontroller (Fig. 1f). The field of view was targeted to the caudal part of Dc (cDc; Fig. 7a). Using a different VR system, neurons in this area were previously found to be active at distinct positions within a virtual corridor containing multiple images as landmarks (Palacios-Flores et al., 2026). Within a typical field of view, hundreds of individual somata could be resolved (Fig. 7b). We extracted time series of relative fluorescence changes (ΔF/F) from regions of interest (ROIs) corresponding to individual somata. To identify spatially selective neurons we defined spatial kernels corresponding to 50 cm space bins and used them as regressors to identify neurons that are selectively active at the corresponding locations. Fig. 7b,c shows activity traces of five neurons that were highly correlated to a given kernel (r = 0.51–0.58; Fig. 7b,c, Methods). Over a period of 44.5 min, the fish traversed the corridor multiple times. Throughout this time, neurons were active with high selectivity when the fish was at the location specified by the kernel (Fig. 7c). These results show that the VR platform supports stable multiphoton calcium imaging during behaviour, allowing for the detection and analysis of position-related activity in cDc (Palacios-Flores et al., 2026).

**Figure 7.**
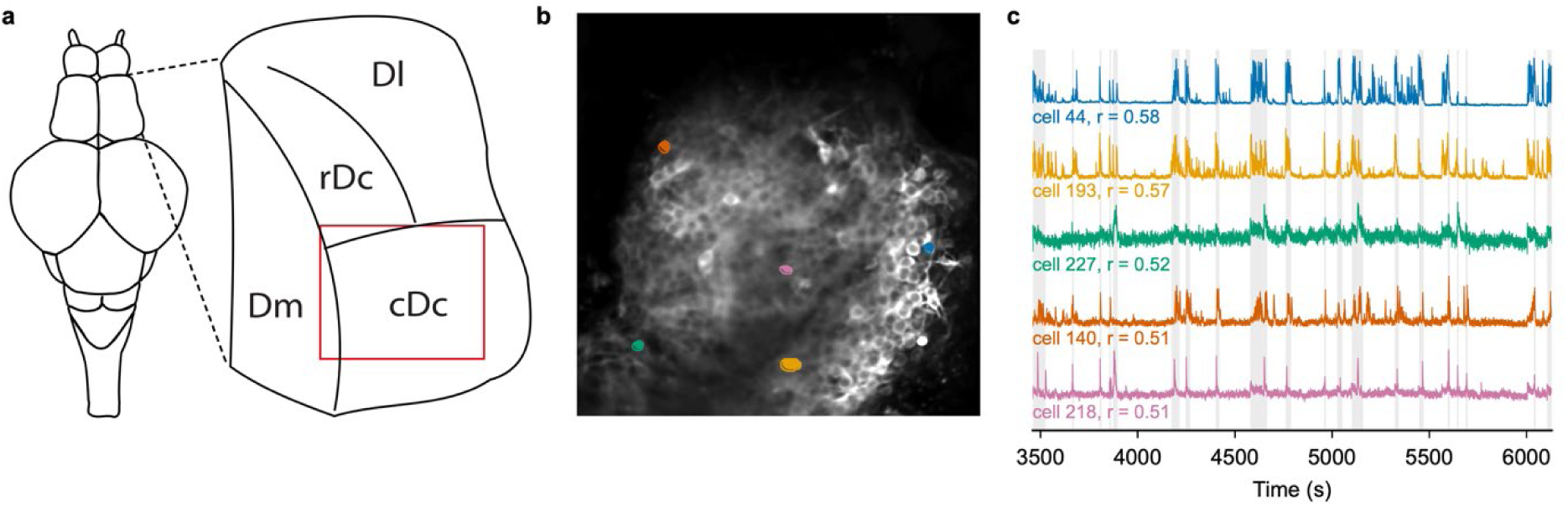
Two-photon calcium imaging of the telencephalon during virtual navigation. **a**, Overview of dorsal telencephalon and region selected for 2-photon imaging (red rectangle). Left: dorsal view of the adult zebrafish brain. Right: enlargement of the right telencephalic hemisphere, showing the medial (Dm), lateral (Dl) and central (rostral, rDc; caudal, cDc) subdivisions of the dorsal pallium. **b**, Raw fluorescence image of cDc in a Tg(neurod:GCaMP6f) fish. 242 cells were segmented in this plane. **c**, Normalized ΔF/F of five cells (coloured as in b) over 44 min of one session. X-axis shows time from session start. Mask depicted by grey shading shows times when the fish was in a specified 50 cm segment of the corridor. Neurons were selected by high correlation between the ΔF/F trace and the mask. Labels show the correlation coefficient (r) and each cell’s index.

## 3 Discussion

We present a versatile platform for VR experiments in head-fixed adult zebrafish that allows for joint analyses of neuronal activity and behavior. The platform comprises hardware and software for head fixation, closed-loop VR operation and the modular design of experimental workflows. In comparison to a previous VR system (Huang et al., 2020), improvements include simplified software, optimized closed-loop control using a microcontroller, enhanced compatibility with commercially available projectors, additional options to trigger external events and to read out behavioral parameters, a new algorithm to predict locomotion from tail videos, more extensive logging of metadata, software for the flexible design of experimental workflows, and hardware options for odor stimulation. Using this platform we found that a visual landmark (stone arch) locally modified swimming behavior of head-fixed adult zebrafish traversing a linear virtual corridor. Moreover, 2-photon calcium imaging results confirmed spatial selectivity of neurons in telencephalic area cDc. These results demonstrate the functionality of the VR platform and illustrate its versatility.

### 3.1 Technical features and functionality

Following previous work (Torigoe et al., 2021; Bandonil et al., 2023; Islam et al., 2024) we used a headplate rather than pins for head fixation, which reduced the complexity and duration of surgical procedures. Hardware and software changes were made in the control loop for VR updates. Tail tracking, rendering, and timing were separated into distinct processes that communicated via the microcontroller. These changes substantially simplified VR implementation and increased the reliability of data logging.

A high quality of VR updates is critical because updates producing unexpected changes in the visual scenery tend to produce non-naturalistic motor outputs that interfere with experimental goals. We therefore designed tail tracking algorithms and reliable procedures to produce high-quality predictions and closed-loop updates. We assessed swimming behavior using three categories: routing swimming, inactivity, and burst swimming. Non-naturalistic swimming is contained primarily in the burst swimming category, along with other rare locomotory patterns. Overall, fish spent ∼96% of their time in the routine swimming or inactive state, which resemble behavioral patterns observed in the home tank, while burst swimming was rare. We therefore conclude that algorithms for tail tracking and locomotor predictions resulted in largely naturalistic swimming in the VR.

Detailed logging of metadata provided new functionality including the readout of additional behavioral information. Stored signals for translational VR updates provide basic information about swimming speed as a function of position and time. In addition, we took advantage of tail shape data to extract information about body posture. Since all data are logged to individual frames, kinematic and postural analyses could be performed with high temporal precision, which enabled detailed analyses of behavioral responses to visual objects.

In a subset of experiments, landmarks in distal parts of the corridor were invisible at the start of the trial and became visible only at a defined distance relative to the fish, thus simulating “murky water”. By varying the position of the landmark in the corridor we then asked whether responses to a landmark depended on visual input, on positional information derived by path integration, or both. This setting may be particularly useful to dissect neuronal mechanisms underlying navigation and further illustrates the versatility of the VR platform.

### 3.2 Applications of the VR platform

To examine behavioral responses to the VR we added a stone arch to the corridor in a subset of trial blocks. The arch had no effects on general swimming behavior but distinct changes in body posture or speed were observed as fish approached the arch. When the arch was visible from a distance, the cumulative tail angle and tail complexity increased near the arch without a concomitant change in swimming speed. We interpret these changes as potential attempts to turn, consistent with observations indicating that adult zebrafish avoid crossing through narrow apertures. When the arch appeared rapidly out of “murky water”, the most pronounced behavioral response was a decrease in swimming speed, indicating that fish performed different maneuvers. These observations illustate how the information accessible through the VR system can be used for granular analyses of swimming behavior.

Observations made after adding or removing an arch in a familiar VR indicate that behavioral responses to landmarks are not stationary but modulated by experience and novelty. While additional experiments are required to further characterize experience-dependent modulation of behavior, the results illustrate how the VR platform can be leveraged to analyze questions related to navigation and spatial memory. VR experiments to address such questions may performed using freely swimming or head-fixed fish (Ahrens et al., 2013; Stowers et al., 2017; Huang et al., 2020; Torigoe et al., 2021; Bandonil et al., 2023; Islam et al., 2024; Demarchi et al., 2025; Sanchez et al., 2025; Tsao et al., 2025; Mills et al., 2026; Palacios-Flores et al., 2026). Our system is designed for head-fixed adult zebrafish to enable joint measurements of behavior and neuronal activity at stages when social and cognitive behaviors are well-developed. Taking advantage of new functionalities for gating of photomultipliers and projectors we performed 2-photon calcium imaging of neuronal activity in the VR and confirmed the presence of spatially selective neurons in telencephalic area cDc (Palacios-Flores et al., 2026). Conceivably, our VR approach should be adaptable to other experimental needs and species including *Danionella* or larger fish. All software and drawings of custom parts are available in public repositories (see Methods).

## 4 Methods

### 4.1 Experimental subjects

Experiments were performed in adult (6 – 18 months old) Tg(neuroD:GCaMP6f) zebrafish (Danio rerio) (Rupprecht et al., 2016) of either sex in a nacre background. Animals were raised and kept under standard laboratory conditions (26 – 27 °C, 13 h/11 h light/dark cycle). All animal procedures were approved by the Veterinary Department of the Canton Basel-Stadt, Switzerland.

### 4.2 Surgical preparation and head fixation

For head fixation, fish were anaesthetised individually in tricaine methanesulfonate (MS-222; 0.24 g/L) until unresponsive to a light tail pinch (<2min) and placed in a custom-built gyroscopic holder. A cannula was inserted into the mouth and anaesthesia was maintained by constant perfusion of the gills with MS-222 (0.074 g/L; flow rate, 4 ml/s). Correct adjustment of the depth of anesthesia is critical. In a subset of experiments, fish additionally received lidocaine (5 mg/L) by bath application while swimming freely. No obvious effect of lidocaine was detected during or after anesthesia, as assessed by recording visual observations of behavioral parameters (rate of opercular movement, swimming speed, swimming pattern, preferred height in the water column, and tail fin position [wafting or lifting]) in scoresheets. MS-222 and Lidocaine were dissolved in system water at pH 6.5– 7.

The skin over the dorsal cranium was removed to expose the skull over the telencephalon, optic tectum and the triangular area on either side of the cerebellum above the gills. A custom 3D printed stainless steel headplate was bonded to the skull with dental cement (GC Fuji 1) caudal to the telencephalon. The cement adhered the headplate to the exposed bone over the caudal end of the optic tectum and to the triangular bones on either side (Fig. 1a-d). This procedure leaves an exposed window over the telencephalon for two-photon imaging. Two C-hooks on either ends of the headplate (radius 3 mm, 22.5 mm apart; CAD drawings: https://github.com/fmi-basel/tailtracking) were used to mount the fish onto the two posts of the tank holder by 2.5 mm screws. (Fig. 1c,d). The body and tail were free to move. The holder with posts was then inserted into a fixed position inside the VR tank filled with system water and the closed-loop visual feedback was started immediately. The time from induction of anesthesia to recovery and swimming in the VR was <25 min.

### 4.3 Virtual-reality apparatus and closed-loop control

The VR tank and projection system were based on a previous design (Huang et al., 2020; Palacios-Flores et al., 2026) with minor modifications. The VR was projected onto diffusive film on the outside of three walls of the semi-hexagonal tank while the fish was positioned approximately at the center of the hexagon (Fig. 1e). The scene was rendered with the Python game engine Panda3D (https://github.com/fmi-basel/fishvr) by three virtual cameras (60° horizontal field of view each, at headings of 0° and ±60°, 180° in total) on a single 5760 × 1080 pixel window, split into one tile per projector. The rendering loop ran at 60 Hz nominally (median frame period 11.11 ms, 99^th^ percentile 18.5 ms). Hardware- and setup-specific configurations such as the geometry of the screens, camera headings and loop rate are read from a configuration file. The VR model was scaled such that one unit distance in VR space corresponds to 1 m in the real world in terms of angular size of objects from the perspective of the fish. The corresponding scaling factor for our setup was 0.72, which is encoded in the configuration file and applied to the 3D model of the VR world.

The tail was illuminated from the side using infrared LEDs and imaged from below through the transparent tank floor with a camera (XIMEA MQ022RG-CM-S7 USB camera; Fig. 1e, right), using a 300 × 360 pixel region of interest centered on the tail, at 150 frames s⁻¹ (6.0 ms exposure). A separate Python process (https://github.com/fmi-basel/tailtracking) fitted seven evenly spaced points on the midline along the tail in real time. Starting from a user-defined coordinate at the base of the tail, the points were placed by identifying the location of the brightest point on the filtered intensity profile of the cross section of the tail at a fixed distance from the previous point (Randlett, 2023). The output was broadcast as a UDP message which is read asynchronously by the VR script.

The change in heading direction was calculated using the cumulative angle between the four most caudal points on the tail, which was low-pass filtered (400 ms time constant) and scaled by a gain factor (gain_h). Because heading is locked in the one-dimensional corridor, it did not turn the scene and served only as a read-out of lateralized tail activity in the present study. gain_h was identical (7.4131 × 10⁻⁴) in all sessions.

Velocity was estimated using the amplitude of oscillations of the tail on either side of the midline. The midline position was computed as the median position of the tail when at rest from the most recent frames over a 15 second window. The peak amplitude of tail-flicks on either side of the midline was extracted over a history of 400 ms. Twice the lower value of the two peak amplitudes was then computed and low-pass filtered with a 400 ms time constant. In a subset of experiments (“murky water”), the velocity estimate was weighted down by turn amplitude to suppress forward displacement while turning. This was achieved by mapping the magnitude of heading change onto a predetermined logistic function ranging from 0 to 1. The velocity estimate was then multiplied by this logistic weight. Finally, the velocity estimate was scaled by a gain factor (gain_v) and soft-clamped below a maximum value of 0.30 m s⁻¹ (clamp absent in two sessions). gain_v was adjusted for each fish at the start of the session with the closed loop running (Huang et al., 2020; Palacios-Flores et al., 2026) so that each animal traversed the corridor with realistic swim speeds and a naturalistic swim pattern. Across fish in this study, gain_v settings took 17 different values that varied over a ∼4.6-fold range.

A Teensy 4.0 microcontroller on a custom interface board synchronised the virtual reality with peripheral hardware. The state of the microcontroller was read and updated on each iteration of the main rendering loop of the VR using an adaptation of the protocol (Desai et al., 2017). The microcontroller was set up to issue two-photon start and end triggers and read the two-photon frame clock and acquisition state. In addition, it drove up to eight stimulation trigger outputs and one analogue voltage output, which can be programmed as higher level parameters in the trial configurator GUI.

A frame-by-frame log was saved as a csv file from both the tail tracking and VR instances. The frame counter value can be used to align the tail tracking log with the corresponding VR frame. In addition, the VR instance also generated a trial-level log of metadata in a JSON file which is used to extract relevant trials and their properties for analysis. A detailed description of the fields in the log files can be found at https://github.com/fmi-basel/tailtracking.

### 4.4 Virtual environment, paradigms and dataset

The environment was a one-dimensional, forward-only corridor with naturalistic textures (rocks, pebbles) on the floor and walls. On a subset of trials, a single stone arch (landmark) was added at a specific position. When the fish reached the end of the corridor, lights were briefly dimmed and the fish was teleported back to the start. Trials were capped at 60 s and incomplete trials were repeated.

The dataset comprises 46 sessions from 34 animals. In each session, one experimental paradigm was applied. Ten animals contributed more than one session. Session identifiers (e.g. 20260108_1b2) encode date, fish, set-up and session number.

### 4.5 Behavioural state classification

Each session was scored for behavioural state from the tail video in 2,000 windows of 45 frames (0.3 s at 150 Hz), sampled across the session. The motion index of a window is the mean absolute frame-to-frame change in grey value, averaged over the pixels brighter than 25 grey levels in either frame of a pair and over the 44 frame pairs of the window (unit: grey levels per frame). A window was scored as active when its motion index exceeded 0.96 grey levels per frame. This threshold was determined empirically by human inspection of video data. An active window was further scored as burst-swimming when the median tail-bend index over its frames was at least 0.30. The bend index is √(L2/L1), where L2/L1 is the ratio of the minor to the major-axis spread of the segmented tail in a principal component analysis; it is near 0 for a straight tail and increases as the tail bends. A fish was classified as inactive if fewer than 20% of its windows were active, as burst-swimming if more than 5% were burst-swimming, and as swimming routinely otherwise. We note that the routine swimming and inactive states were classified consistently by humans and resembled naturalistic behaviors in the home tank. The burst-swimming state was more heterogeneous, containing events that resembled struggling but also movements that were interpreted as attempted turns or other maneuvers involving large tail bends. Since burst-swimming was rare, we did not attempt to further sub-classify this category.

### 4.6 Data processing and trial inclusion

Within each session, data were split into trials and each trial was divided into 140 positional bins. The final 10% of corridor bins were excluded from analysis because virtual speed typically changed at the corridor end. Tail-camera samples were aligned to the VR timeline through the hardware frame counter.

Only trials in which the animal covered at least 80% of the corridor were analyzed (22–43% of trials depending on paradigm, with no bias by condition, p >0.3). Trials were averaged within sessions and sessions within animals before any group statistic. Statistical comparisons were thus performed between animals.

### 4.7 Landmark-based analyses

Because landmark positions differed between animals and trials, most analyses were aligned to each trial’s landmark position. In “Landmark addition” and “Landmark deletion” paradigms (Fig. 5), cumulative tail angle was analyzed using each animal’s first block switch only (10 completed trials before and after landmark switch). The reported contrast is a double difference: (Landmark present) – (Landmark absent) in the landmark region minus the same difference over the rest of the corridor. The landmark region was defined as the space bin containing the landmark and the two adjacent space bins on either side (5 bins total). Analyses of tail complexity used all switches, with trial offsets counted over completed trials.

For displacement (Fig. 6), panels a-d use the same first-transition window and panels e,f use all data. For statistical comparisons, all within-session (Landmark present) – (Landmark absent) differences were used. Anticipatory slowing was quantified because it is has been used as a behavioural measure in rodent virtual corridors (Youngstrom and Strowbridge, 2012; Sato et al., 2017). The onset of slowing was estimated based on the speed difference against distance from the landmark by uncorrected per-bin t-tests (descriptive only). Pre-crossing speed (1 s before the crossing) was compared between (Landmark present) and (Landmark absent) trials at the same corridor location, paired within animals, and transit time from first sight (0.5 m) to the landmark was measured from the logged trajectories.

### 4.8 Two-photon imaging

Calcium imaging was performed using a custom-built two-photon microscope equipped with a Ti:sapphire laser (MaiTai DeepSee, Spectra-Physics), 8 kHz resonance scanners, gated photomultiplier tubes (H11706P-40, Hamamatsu), a 16× water-immersion objective (Nikon N16XLWD-PF) (Huang et al., 2020), and ScanImage software (Pologruto et al., 2003). Single-plane images of 512 × 512 pixels, covering a field of view of approximately 255 × 255 µm, were acquired at 30 Hz approximately 45 µm below the dorsal surface of the telencephalon in Tg(neurod:GCaMP6f) fish. The microcontroller logged the ScanImage frame clock together with the behavioural data on the same clock, so that every imaging frame could be assigned to the position of the fish in the virtual corridor at the time it was acquired (Fig. 1e). Image frames were motion-corrected by two rounds of phase correlation against the average of the stack. Regions of interest corresponding to individual neurons were extracted with Cellpose-SAM (Pachitariu et al., 2025) and curated manually. Fluorescence was expressed as ΔF/F, with F0 defined as the 10th percentile of each cell’s fluorescence distribution. For Fig. 7c, a binary regressor marking the frames in which one fish over one session occupied one third of the corridor was correlated with the ΔF/F trace of every cell, and the five cells with the highest correlation values are shown.

### 4.9 Statistics

All tests were two-sided and computed across animals. One-sample t-tests against zero were used for within-animal differences (turning contrast, curvature response, anticipatory speed change). Paired t-tests were used for present-versus-absent comparisons at matched positions and for first-versus-later appearances. Cluster-based permutation testing was performed to analyze the spatially extended speed difference. Exact p values, effect sizes and tests are given in the legends (* p < 0.05, ** p < 0.01, *** p < 0.001).

### 4.10 Software, code and data availability

Analysis were performed in Python. The virtual-reality engine, tail tracker, Teensy firmware and board design, configuration files, corridor models are provided as companion repositories:

https://github.com/fmi-basel/tailtracking

https://github.com/fmi-basel/fishvr

## Acknowledgements

We thank Markus Klement for expert mechanical service, the FMI fish facility for fish care, Estelle Arn for lab management, and the Friedrich lab for fruitful discussions. This work was supported by the Novartis Research Foundation, by the European Research Council (ERC; grant agreement no. 101167289), by the the Swiss National Science Foundation (grants no. 310030_212236, PZ00-3_232850), and by EMBO postdoctoral fellowships to S.N. and L.A. (ALTF 1060-2024, ALTF 379-2022).

## References

Ahrens MB, Huang K-H, Narayan S, Mensh BD, Engert F (2013) Two-photon calcium imaging during fictive navigation in virtual environments. Front Neural Circuits 7:104.

Ahrens MB, Li JM, Orger MB, Robson DN, Schier AF, Engert F, Portugues R (2012) Brain-wide neuronal dynamics during motor adaptation in zebrafish. Nature 485:471–477.

Bandonil JS, Liao Y-H, Fathi A, Huang K-H (2023) Two-Photon Calcium Imaging of Forebrain Activity in Behaving Adult Zebrafish. Journal of Visualized Experiments (JoVE):e65526.

Bianco IH, Kampff AR, Engert F (2011) Prey capture behavior evoked by simple visual stimuli in larval zebrafish. Front Syst Neurosci 5:101.

Bloch S, Froc C, Pontiggia A, Yamamoto K (2019) Existence of working memory in teleosts: Establishment of the delayed matching-to-sample task in adult zebrafish. Behavioural Brain Research 370:111924.

Chou M-Y, Amo R, Kinoshita M, Cherng B-W, Shimazaki H, Agetsuma M, Shiraki T, Aoki T, Takahoko M, Yamazaki M, Higashijima S, Okamoto H (2016) Social conflict resolution regulated by two dorsal habenular subregions in zebrafish. Science 352:87–90.

Demarchi L, Coraggioso M, Hubert A, Panier T, Morvan-Dubois G, Bormuth V, Debrégeas G (2025) Logarithmic coding leads to adaptive stabilization in the presence of sensorimotor delays. Proceedings of the National Academy of Sciences 122:e2510385122.

Desai NS, Gray R, Johnston D (2017) A Dynamic Clamp on Every Rig. eNeuro 4 Available at: https://www.eneuro.org/content/4/5/ENEURO.0250-17.2017.

Friedrich RW, Jacobson GA, Zhu P (2010) Circuit Neuroscience in Zebrafish. Current Biology 20:R371–R381.

Fry SN, Rohrseitz N, Straw AD, Dickinson MH (2008) TrackFly: Virtual reality for a behavioral system analysis in free-flying fruit flies. Journal of Neuroscience Methods 171:110–117.

Gerlai R (2017) Zebrafish and relational memory: Could a simple fish be useful for the analysis of biological mechanisms of complex vertebrate learning? Behavioural Processes 141:242–250.

Harvey CD, Collman F, Dombeck DA, Tank DW (2009) Intracellular dynamics of hippocampal place cells during virtual navigation. Nature 461:941–946.

Heisenberg M, Wolf R (1979) On the fine structure of yaw torque in visual flight orientation ofDrosophila melanogaster. J Comp Physiol 130:113–130.

Huang K-H, P. Rupprecht, Frank T, Kawakami K, Bouwmeester T, Friedrich RW (2020) A virtual reality system to analyze neural activity and behaviour in adult zebrafish. Nature Methods 17:343–351.

Islam T, Torigoe M, Tanimoto Y, Okamoto H (2024) Adult zebrafish can learn Morris water maze-like tasks in a two-dimensional virtual reality system. Cell Reports Methods 4:100863.

Jouary A, Haudrechy M, Candelier R, Sumbre G (2016) A 2D virtual reality system for visual goal-driven navigation in zebrafish larvae. Sci Rep 6:34015.

Kentros CG, Agnihotri NT, Streater S, Hawkins RD, Kandel ER (2004) Increased attention to spatial context increases both place field stability and spatial memory. Neuron 42:283–295.

Lindeyer CM, Reader SM (2010) Social learning of escape routes in zebrafish and the stability of behavioural traditions. Animal Behaviour 79:827–834.

Mills SC, Beldade R, Lafaye J, Lecchini D, Vidal M (2026) A fully immersive Virtual Reality setup using DeepLabCut reveals species-specific behavioural responses of freely moving coral reef fishes to conspecifics and predators. HAL open science.

O’Keefe J, Dostrovsky J (1971) The hippocampus as a spatial map. Preliminary evidence from unit activity in the freely-moving rat. Brain Res 34:171–175.

Oliveira RF (2013) Mind the fish: zebrafish as a model in cognitive social neuroscience. Front Neural Circuits 7:131.

Orger MB, Kampff AR, Severi KE, Bollmann JH, Engert F (2008) Control of visually guided behavior by distinct populations of spinal projection neurons. Nat Neurosci 11:327–333.

Ortiz C, Allegra M, Schmidt-Hieber C (2025) Task engagement differentially drives hippocampal and neocortical neural codes. :2025.06.26.661816 Available at: https://www.biorxiv.org/content/10.1101/2025.06.26.661816v1.

Pachitariu M, Rariden M, Stringer C (2025) Cellpose-SAM: superhuman generalization for cellular segmentation. :2025.04.28.651001 Available at: https://www.biorxiv.org/content/10.1101/2025.04.28.651001v1.

Palacios-Flores K, Eckhardt J, Huang K-H, Narayanan S, Friedrich RW (2026) Dynamic cognitive representations in the dorsal pallium of adult zebrafish. Available at: https://www.biorxiv.org/content/10.1101/2025.05.20.655100v2.

Pettit NL, Yuan XC, Harvey CD (2022) Hippocampal place codes are gated by behavioral engagement. Nature Neuroscience 25:561–566.

Pologruto TA, Sabatini BL, Svoboda K (2003) ScanImage: Flexible software for operating laser scanning microscopes. BioMedical Engineering OnLine 2:13.

Radvansky BA, Dombeck DA (2018) An olfactory virtual reality system for mice. Nat Commun 9:839.

Randlett O (2023) pi_tailtrack: A compact, inexpensive and open-source behaviour-tracking system for head-restrained zebrafish. J Exp Biol 226:jeb246335.

Rodríguez F, Quintero B, Amores L, Madrid D, Salas-Peña C, Salas C (2021) Spatial Cognition in Teleost Fish: Strategies and Mechanisms. Animals 11:2271.

Rupprecht P, Prendergast A, Wyart C, Friedrich RW (2016) Remote z-scanning with a macroscopic voice coil motor for fast 3D multiphoton laser scanning microscopy. Biomed Opt Express, BOE 7:1656–1671.

Sanchez S, Escobedo R, Bastien R, Lenseigne B, Denis A, Moreau M, Combe M, Straw AD, Sire C, Theraulaz G (2025) An open-source closed-loop Virtual Reality system to investigate social interactions and collective behavior in fish. :2025.05.20.655100 Available at: https://www.biorxiv.org/content/10.1101/2025.05.20.655100v2.

Sato M, Kawano, M., Mizuta, K., Islam, T., Lee, M. G., Hayashi, Y. (2017) Hippocampus-dependent goal localization by head-fixed mice in virtual reality. eNeuro 4; 10.1523/ENEURO.0369-16.2017.

Schluessel V, Kreuter N, Gosemann IM, Schmidt E (2022) Cichlids and stingrays can add and subtract ‘one’ in the number space from one to five. Sci Rep 12:3894.

Stowers JR, Hofbauer M, Bastien R, Griessner J, Higgins P, Farooqui S, Fischer RM, Nowikovsky K, Haubensak W, Couzin ID, Tessmar-Raible K, Straw AD (2017) Virtual reality for freely moving animals. Nat Methods 14:995–1002.

Torigoe M, Islam T, Kakinuma H, Fung CCA, Isomura T, Shimazaki H, Aoki T, Fukai T, Okamoto H (2021) Zebrafish capable of generating future state prediction error show improved active avoidance behavior in virtual reality. Nat Commun 12:5712.

Tsao J-Y, Tang C-Y, Chen C-Y, Yang C-T, Huang K-H (2025) Integration of social and depth information during affiliative behavior in zebrafish. Current Biology 35:3429–3439.e4.

Yang C, Mammen L, Kim B, Li M, Robson D, Li JM (2024) A population code for spatial representation in the zebrafish telencephalon. Nature 634:397–406.

Yang E, Zwart MF, James B, Rubinov M, Wei Z, Narayan S, Vladimirov N, Mensh BD, Fitzgerald JE, Ahrens MB (2022) A brainstem integrator for self-location memory and positional homeostasis in zebrafish. Cell 185:5011–5027.e20.

Youngstrom IA, Strowbridge BW (2012) Visual landmarks facilitate rodent spatial navigation in virtual reality environments. Learning & Memory 19:84–90.

